# Using a Motion Sensor to Categorize Low Back Pain Patients: A Machine Learning Approach

**DOI:** 10.1101/803155

**Authors:** Masoud Abdollahi, Sajad Ashouri, Mohsen Abedi, Nasibeh Azadeh-Fard, Mohamad Parnianpour, Ehsan Rashedi

## Abstract

Low back pain (LBP) remains a critical health issue impacting literally millions of people worldwide. Currently, clinical practitioners rely on subjective measures such as the STarT Back Screening Tool to categorize LBP patients, which then informs specific treatment regimens. This study sought to develop a machine learning model to classify LBP patients into different groups according to kinematic data. Specifically, an inertial measurement unit (IMU) was attached to each patient’s chest while he performed trunk flexion/extension motions at a self-selected pace. Machine learning algorithms such as support vector machine (SVM) and multi-layer perceptron (MLP) were implemented to evaluate the efficiency of the models. The results showed that the kinematic data we obtained could be used to categorize the patients into two groups: high vs. low-medium risk. We achieved accuracy levels of ~75% and 60% for SVM and MLP, respectively. Additionally, among a range of variables detailed herein, we determined that time-scaled IMU signal resulted in the highest accuracy. Our findings support the use of body-motion measures in developing prognosis tools for healthcare applications. Our results could help overcome the need for objective clinic-based diagnosis approaches, which in turn would lead to assigning better treatment approaches and rehabilitation services for LBP sufferers.

## Introduction

Low back pain (LBP) remains a challenging and often debilitating health condition that is considered to be the most common cause of disability and the most widespread musculoskeletal disorder worldwide [1, 2]. It is estimated that around 84 percent of the adult population will experience LBP at some time in their lives, and up to 33% of adults may be dealing with this condition at any given moment in time [3–5]. In both developed/developing countries, LBP has been identified as the fourth-most frequent condition warranting a visit to a physician, the fifth-most common reason for hospitalization, and the third-most frequent cause for undergoing a surgical operation [4, 6]. There is, therefore, a compelling rationale for investigating this widespread condition in order to identify more appropriate treatments and injury-prevention approaches.

Meanwhile, researchers have verified that there is no one-size-fits-all treatment approach for LBP patients given the complexity of condition and the heterogeneity of LBP causes among sufferers [7, 8]. However, evidence-based guidelines do confirm the necessity of stratifying LBP patients in primary care according to (a) their risk for subsequent disability, and (b) specific treatment approaches for each group [9, 10]. While it may seem straightforward to partition treatment groups in this way, it it by no means easy to assign each patient to the most suitable LBP group/treatment scenario. To address this dilemma, several methods have been developed to classify LBP patients [11–14]. The most commonly accepted instrument for stratifying treatment based on prognosis risk is the STarT Back Screening Tool (SBST) [11, 15–18]. This approach requires LBP patients to complete a screening questionnaire, whose results are used to assign the respondent into one of three groups: low, medium, and high risk [19]. For each of the three LBP groups, there exists a specific treatment approach. Notably, one of them, i.e., the high-risk group, requires the patients to undergo a series of six individual physiotherapy sessions over three months. In contrast, the low- and medium-risk groups are treated with less aggressive approaches, such as providing therapeutic information to the patients. Hence, it is important to distinguish the high-risk individuals among these patients for much-needed curative therapies.

To categorize LBP patients for appropriate treatment protocols, clinical decision-making in a primary care environment is currently based on qualitative questionnaires, and thus the pitfalls of subjective reporting. In other words, although stratifying patients into different groups and assigning different treatment approaches has led to better therapeutic outcomes (e.g., decreasing disability from LBP and reducing the time off work), this classification approach tends to be prone to error due to the low reliability of the subjective information collected from patients [20]. In contrast, advancements in sensor technology are leading to efforts to augment subjective measures with objective approaches, such as measuring patient movements during walking or other controlled tasks [21–25]. The core concept of these studies involves utilizing differences in trunk motion (i.e., trunk kinematics) for people suffering from LBP.

Researchers continue to investigate kinematic approaches for their diagnostic potential, or even as a tool to make a clinical decision in primary care [21, 26–31]. Some earlier studies involving LBP sufferers have confirmed a correlation between the quality of motion in people and their health status [32–35]. Marras et al. implemented several models to classify healthy versus LBP people based on trunk angular motion features during flexion/extension and bending of the trunk in various symmetric and asymmetric planes of motion [33, 36]. Similarly, Ashouri et al. was able to distinguish healthy people from LBP patients based on a signal from an inertial measurement unit (IMU) sensor on the chest during a trunk flexion/extension task using a Support Vector Machine (SVM) classifier [21]. In addition to focusing on LBP, similar studies have focused on the neck region. For example, Bahat et al. revealed that neck pain would lead to lower peak and mean velocity of the neck during flexion/extension motions [37].

All of these prior studies sought to investigate a model that would distinguish healthy subjects from a patient cohort. In contrast, there are far fewer scholarly reports designed to stratify LBP patients and correlate those findings with proper treatment approaches. This study, therefore, was designed to categorize LBP sufferers using a new objective method. Specifically, we utilized an IMU sensor placed on the torsos of LBP patients to collect kinematic data. We then utilized different machine learning approaches to classify our study subjects into different subgroups based on trunk kinematics, and then compared the results of these data analytics approaches to identify the most accurate one. This study is expected to contribute to the ability to objectively diagnose and categorize LBP according to severity, which will help clinicians and care providers make more accurate decisions about the risk levels for individuals with LBP.

## Materials and Methods

A system diagram is provided to depict the overall procedures for data collection and data analysis in this study (Fig 1). Details about each aspect of the study are provided in the following sections.

**Figure 1.**
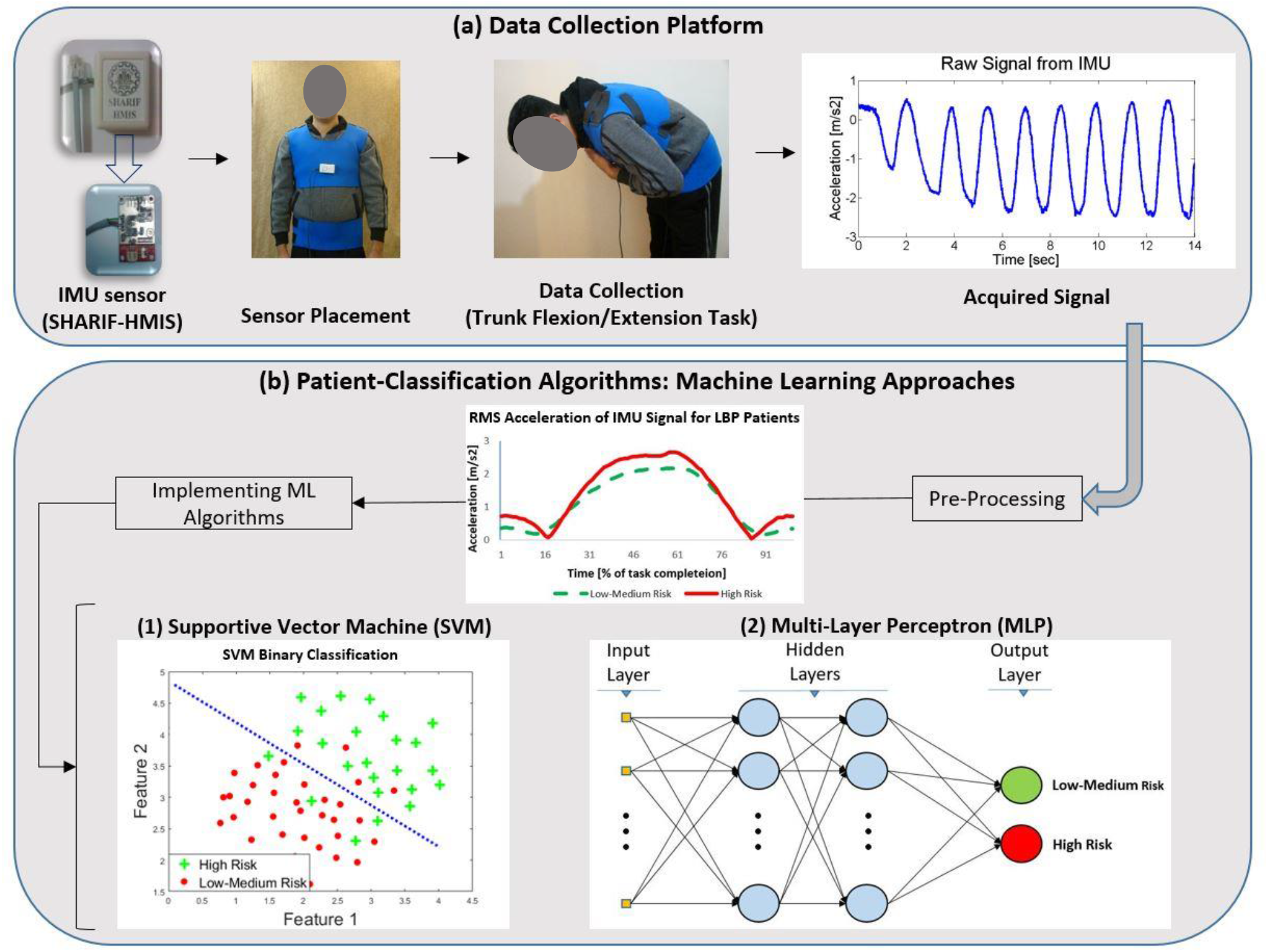
Schematic system diagram of the platform for the classification of LBP patients. (a) Sensor and data collection procedure. (b) Data analysis procedure.

### Participants

Ninety-four male volunteer LBP patients were recruited for this study. The average (standard deviation) age, height, and weight of the participants were 43.6 (6.9) years, 172.6 (7.3) cm, and 79.5 (12.5) kg, respectively. Statistical analyses, i.e., ANOVA and t-test, were conducted to ensure that there were no significant differences (p-value > 0.05) among different groups in terms of age, height, weight, and BMI. Three inclusion criteria were implemented to recruit participants: 1) Participants had to be between 20-50 years of age who identified with low back pain based on Waddell et al.’s definition [38]; 2) During the assay, pain intensity had to remain less than 5 on the Visual Analog Scale (VAS) [39, 40]; and 3) Any participant with a history of spinal surgery was excluded from taking part. All LBP patients were examined by an orthopedic surgeon to meet all the inclusion criteria, and participation was wholly voluntary. Each individual was also required to sign a consent form approved by the Shahid Beheshti University of Medical Sciences Ethics Committee prior to taking part in this study.

### Experimental design

Each subject was asked to perform as many trunk flexion/extensions as he could within a 14-second period. An IMU sensor (9DOF Razor IMU, Sparkfun^®^, Niwot, Colorado) composed of a 3-axis accelerometer and a 3-axis gyroscope with a sampling frequency of 20 Hz was implemented to acquire trunk kinematic data. The sensor was placed on the sternum of the patients during the task (Fig 1) to collect acceleration and angular velocity. In addition to kinematic data, a balance board was used to collect data related to the center of pressure (COP). Four variables were measured from the balance board: x and y range, path length, and area of the ellipse, which captured 95% of the COP data. Moreover, each participant was required to complete the Hospital Anxiety Depression Scale (HADS) [41] and the Tampa Scale of Kinesiophobia (TSK) [42] questionnaires. These quantitative measures were implemented to help verify the objective accuracy of resulting chest kinematic data. Finally, the subjects were asked to fill in the Persian version of the STarT questionnaire [43], which divided (labeled) the LBP patients into three risk groups (low, medium, and high). This data was implemented as the ground truth data for our machine learning algorithms. During the supervised learning algorithm portion of this study, the labeled data was divided into two segments in order to train the model and calculate its sensitivity, specificity, and the accuracy.

### Data processing

For each subject, the collected data were processed in MATLAB (Mathworks, Inc., Natick, MA, USA) to provide a feature vector to be implemented for classification. In this study, there were four performance-oriented feature types: (1) acceleration, (2) velocity, (3) angular acceleration, and (4) force platform data. The first three features included data in X, Y, and Z directions, which were defined as Vertical, Medio-lateral, and Antero-posterior directions, respectively.

The data for each subject comprised several trunk flexion/extension cycles, which could be different depending on the individual. Then, all of the flexion/extension cycles were aggregated into one cycle in order to produce a single profile representing the first three features for each subject. This step led to determining angular/linear acceleration and angular velocity for each subject in only one resultant cycle. Thus, for each subject there was only one acceleration graph, which resulted from averaging (using root mean square) all of the acceleration profiles during flexions/extensions during the 14-second interval. Furthermore, the data were time-scaled such that for each feature profile of the first three features, there were only 100 points. Once the variable of time was removed from the data, the resulting data was the features’ profile in each percentile of task completion. Moreover, 16 statistical features such as max, standard deviation, kurtosis, and skewness, were calculated for each signal.

Since participants were asked to perform flexion/extension as fast as they comfortably could within a specific time (i.e., 14 seconds), the number of cycles differed from person to person. To consider this performance aspect, the angular acceleration of the trunk was also calculated from the angular velocity signal applied in the study. It should be noted that by removing the time variable from the signals through time-scaling and averaging the cycle for each trial, the effect of the different number of cycles for each trial could be eliminated. To shed further light on point, assume two different signals for angular velocity—a single cycle and four cycles. In this instance, the resultant signals would be the same as long as they have the same range and trend of angular velocity. This factor was the primary reason for adding angular acceleration as a study measure (in addition to angular velocity and linear acceleration); angular acceleration would preserve needed data about changes in angular velocity. Apparently, more flexion/extension cycles would result in a larger range of angular acceleration. In other words, the range of angular acceleration corresponds to the number of cycles for each subject.

The last factor incorporated in the analysis represents the four measures obtained from the force platform underneath the participants’ feet: displacement range of center of pressure (COP) in both the x and y directions, COP path length during the task, and ellipse area of the center of pressure amplitude. These features enabled the algorithm to consider balance factors as a part of the discriminating protocol. Finally, prior to applying any machine learning algorithm to the data, all of the features were scaled. All the features implemented for this investigate are listed in Table 1.

**Table 1.**
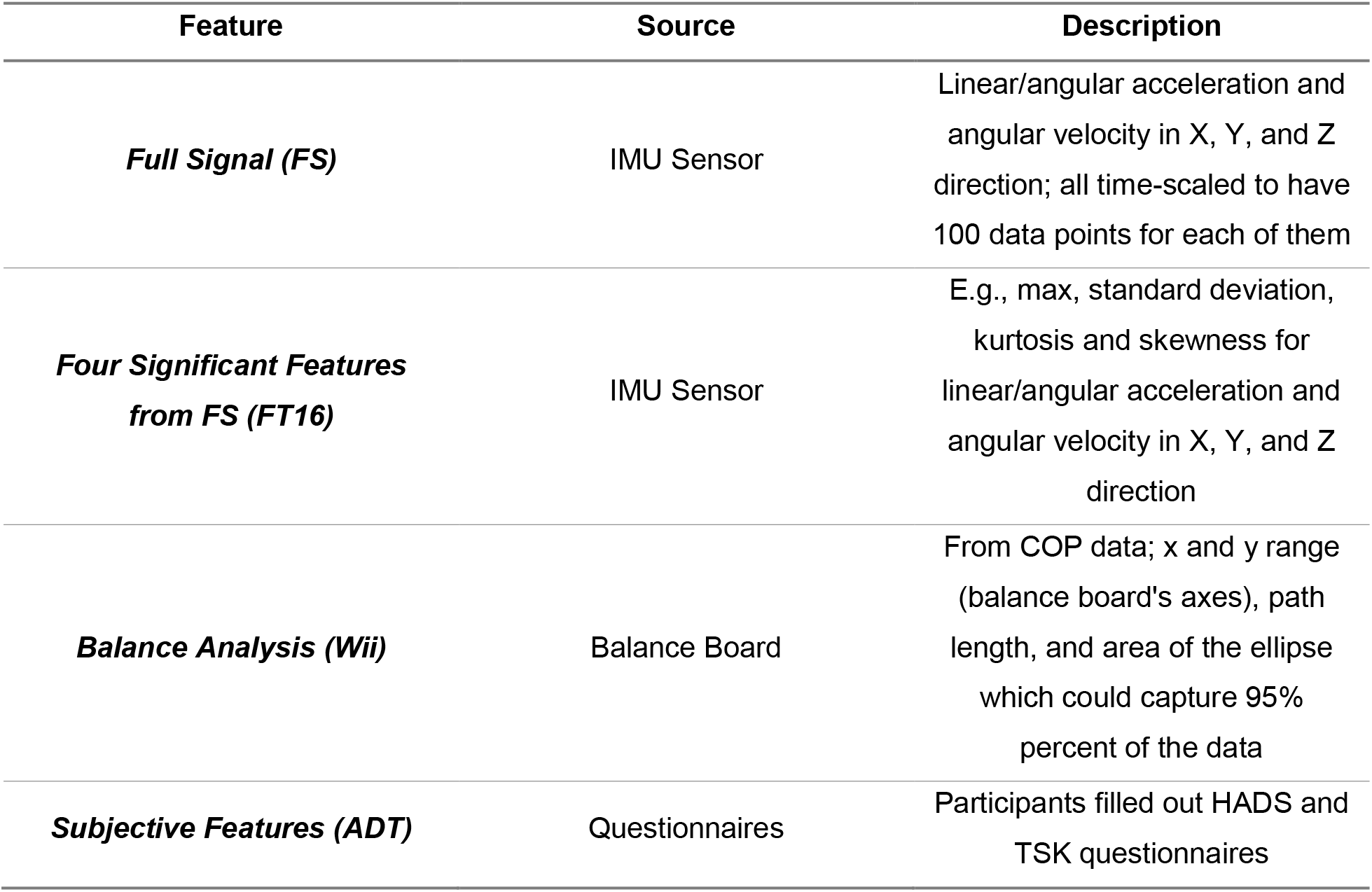
The description of various features and their sources

### Classification approaches

After data processing, three algorithms were implemented in MATLAB software to classify the patients into distinct risk groups. First, the K-means algorithm was implemented to cluster study participants based on their kinematic data and with no use of any labels (unsupervised learning). In the next step, data were evaluated using the support vector machine (SVM) algorithm. Finally, a neural network approach was applied to the data. To determine the best discriminating approach, the feature set of FS (see Table 1) was implemented for all of the algorithms.

### I. K-means

K-means is an unsupervised learning method to cluster data without labeling. In this study, we used K-means to (1) check how many subgroups could appropriately represent all LBP patients based on trunk kinematics, and (2) assess if the kinematic data were properly classified. The latter objective was intended to provide general perceptions about the kinematic data from all patients. The algorithm was initiated from some initial points (equal to the number of clusters), and then added each point to one of these clusters and updated the center points. Finally, all of the sample data were assigned to one of the clusters. The important point that had to be considered was finding the optimal number of clusters, which was essential prior to implementing the K-means algorithm since the number of clusters was the input of the algorithm.

The Calinski-Harabasz index was implemented to determine the proper clustering number. The formulation of the index was as follows [44]:

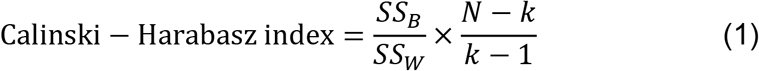

Where *SS_B_* is overall between-cluster variance, *SS_W_* is the overall within-cluster variance, N is the total number of observations, and k is the number of clusters. The number of clusters was determined such that this index could achieve the maximum possible value. The first ratio illustrates the case of having more variance between clusters and less variance in each cluster, thus causing the index to increase. In other words, if the clusters are far away from each other, the index would increase. These data were calculated for the different features, including angular velocity, linear acceleration, and the combination of them.

### II. Support Vector Machine (SVM)

The SVM method was implemented to see how the labeled data could be discriminated using a supervised learning algorithm. For this approach, leave-one-out cross-validation was used, which is useful for finding a hyperplane that can distinguish two groups with high accuracy. The kernel function was selected based on the best outcome among Gaussian, polynomial, and radial basis kernel function. The SVM algorithm was run for each of the subjects such that all other subjects were considered as training data in that particular run. The algorithm predicted the label of the patient using the relevant trained network.

Initially, multi-class SVM was used to segregate patients into the three risk groups: low, medium, and high. Since the accuracy of the three-class SVM approach for all combinations of the features was less than 40%, we decided to combine two groups and compare it to the third one. Toward this end, the SVM algorithm was run to calculate the accuracy, sensitivity, and specificity of the method based on the comparison of the predicted label and ground truth.

For the SVM method, the balance of the dataset could make a significant difference in the accuracy of the discrimination. Since patients were being discriminated into two groups, the sample size for the first group was almost twice the size of the second. To address this concern, some sample data were randomly selected from the larger group so that we ended up with two groups of equal sample size. For example, in the case of two groups of high vs. low-medium risk, in addition to the 28 high-risk patients, 28 patients were randomly selected from the pool of 66 low-medium risk patients. This procedure was performed ten times, and the model was run to avoid the effects of random biased sampling. The same dataset was used for other machine learning tools in order to ensure that we could compare data. The results of the analysis were reported as mean and standard deviation of the ten runs.

### III. Neural Network

Multilayer perceptron (MLP) was implemented for the data. For each subject, there were 900 features (FS feature set), which were considered as the neuron of the input layer. The architecture of the network was considered with eight hidden layers of 700, 500, 300, 100, 50, 15, 10, and 5 neurons. These values were determined based on a manual trial-and-error process to increase the accuracy of the model. The implemented network was a 10-layer feed-forward network. The activation function was a sigmoid function for the hidden layers, and softmax transfer function was applied for the output layer. Eighty percent of the data was utilized in training, and the rest was used to test the model. The algorithm was run for 3-class classification. Since the accuracy for all combination of features was not larger than 40%, similar to the SVM approach we utilized 2-class classification with the same dataset.

As mentioned earlier, all three machine learning algorithms, i.e. SVM and MLP, were run for the feature set of FS. However, the approach leading to the highest accuracy was run for all combination of the features. Accordingly, a full factorial combination of the four feature sets (2^4^ − 1 = 15 cases) was analyzed. A comparison of the accuracy, sensitivity, and specificity for all for these cases produced the feature set(s) with the highest accuracy.

## Results

Analyzing the Calinski-Harabasz index for different features (i.e., angular velocity and linear acceleration) revealed that the maximum value for all the plots (representing different cluster numbers) occurred for the two clusters (Figure 2). In other words, by clustering the data into two groups, we achieved the most significant variance between the clusters, as well as the lowest variance within each cluster. To evaluate the efficiency of k-means clustering, each cluster was labeled based on the largest population of the sample from each group. For example, assume that there were 40 samples in the first cluster in which there were 34 samples with the actual label of ‘A’. In this case, the cluster was labeled as the ‘A’ group, and the other cluster was labeled as the ‘B’ group. Based on this approach, accuracy, sensitivity, and specificity for the K-means algorithm were reported to be 57%, 43%, and 63%, respectively.

**Figure 2.**
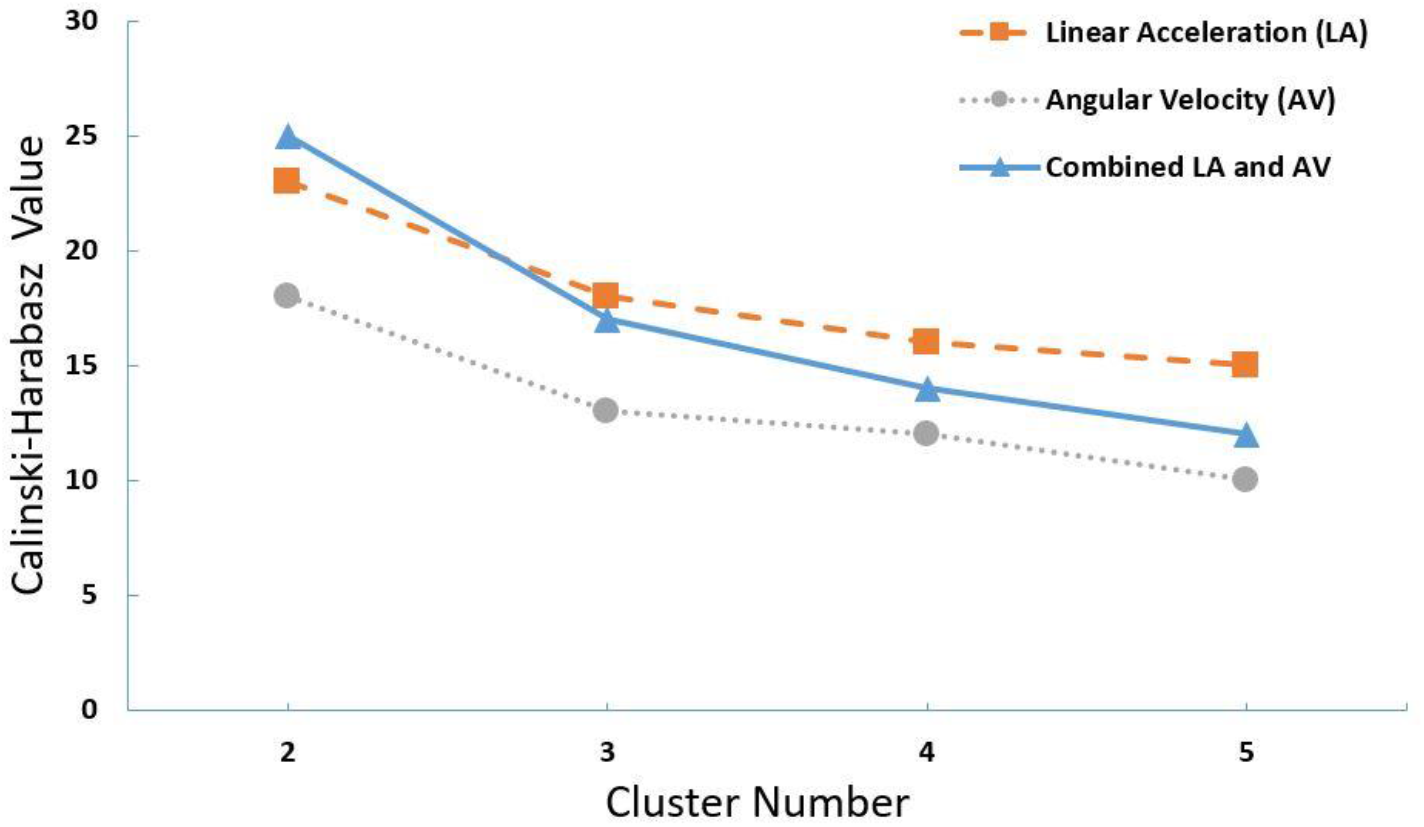
Calinski-Harabasz index for different features including linear acceleration, angular velocity, and the combination of them, for different number of clusters

After implementing the SVM and MLP algorithms for 2-class classification of the data, accuracy, sensitivity, and specificity of the approaches were calculated for all possible 2-class categories (see Table 2). As noted earlier, the sample size for each of the two selected classes was made equal, which required a random selection of samples from the larger group (among the two classes). In order to avoid any biased selection, this process was repeated ten times. Hence, the reported numbers were the mean of ten runs and the relevant standard deviation values (Table 2).

**Table 2.** The accuracy, sensitivity, and specificity of the different combinations of the classes for SVM and MLP algorithms

|  |  | *Low vs.*<br>*Medium-High* | *Medium vs.*<br>*Low-High* | *High vs*<br>*Low-Medium* |
| --- | --- | --- | --- | --- |
| *SVM* | *Accuracy* | 46.3 (6.3) | 45.5 (6.8) | 75.4 (4.2) |
|  | *Sensitivity* | 45.0 (10) | 59.7 (7.5) | 72.5 (3.8) |
|  | *Specificity* | 47.6 (8.6) | 31.4 (7.2) | 78.2 (5.3) |
| *MLP* | *Accuracy* | 50.2 (4.2) | 43.2 (5.6) | 60.2 (5.3) |
|  | *Sensitivity* | 44.7 (10.1) | 43.8 (9.2) | 66.4 (8.1) |
|  | *Specificity* | 55.5 (6.4) | 42.5 (8.6) | 54.0 (9.8) |

Since the highest accuracy was achieved in the SVM approach for discriminating high vs., low-medium risk, the same algorithm was run for the full factorial combination of the feature sets to evaluate the effects of different types of features on the accuracy of the model (Table 3).

**Table 3.** SVM accuracy, sensitivity, specificity for high vs. low-medium classification considering different feature sets including full signal (FS) which was the processed signal of IMU, 16 features from that signal (FT16), the output of balance board (Wii), and finally HADS and TSK questionnaire data (ADT). The reported numbers are in the format of the mean (standard deviation) of the ten runs.

|  | *FS* | *FT16* | *Wii* | *ADT* | *FS*<br>*+FT16* | *FS*<br>*+Wii* | *FS*<br>*+ADT* | *FT16*<br>*+Wii* | *FT16*<br>*+ADT* | *Wii*<br>*+ADT* | *FS*<br>*+FT16*<br>*+Wii* | *FS*<br>*+FT16*<br>*+ADT* | *FS*<br>*+Wii*<br>*+ADT* | *FT16*<br>*+Wii*<br>*+ADT* | *ALL* |
| --- | --- | --- | --- | --- | --- | --- | --- | --- | --- | --- | --- | --- | --- | --- | --- |
| *Accuracy* | 75.4<br>(4.2) | 66.9<br>(5.4) | 39.5<br>(8.3) | 54.8<br>(6.8) | 63.5<br>(5.6) | 65.2<br>(6.6) | 66.6<br>(6.3) | 58.9<br>(5.1) | 52.7<br>(7.4) | 46.8<br>(4.8) | 64.8<br>(5.5) | 66.8<br>(6.2) | 64.2<br>(4.9) | 54.9<br>(6.2) | 59.3<br>(5.4) |
| *Sensitivity* | 72.5<br>(3.8) | 58.3<br>(5.1) | 40.4<br>(8.8) | 55.7<br>(6.9) | 65.8<br>(5.5) | 61.6<br>(7.9) | 67.4<br>(8.7) | 52.1<br>(5.5) | 42.5<br>(8.6) | 48.2<br>(5.3) | 58<br>(4.9) | 68.4<br>(6.7) | 64<br>(6.1) | 44.7<br>(7.7) | 61.5<br>(4.8) |
| *Specificity* | 78.2<br>(5.3) | 75.7<br>(6.5) | 38.6<br>(9.8) | 53.9<br>(7.3) | 61.2<br>(7.3) | 68.7<br>(5.5) | 65.7<br>(7.7) | 63.6<br>(8.9) | 62.8<br>(8.5) | 45.3<br>(6.9) | 71.6<br>(6.4) | 65.1<br>(7.4) | 64.4<br>(5.5) | 65.1<br>(7.6) | 65.4<br>(6.7) |

## Discussion

This study sought to develop a machine learning-based model having the capability of classifying LBP patients into different risk groups (treatment scenarios) based on trunk kinematic data obtained from an IMU sensor on the chest, and then compare that data with results obtained from the STarT questionnaire. Prior information from participants who completed the STarT questionnaire was used to categorize each person as being a high or low-medium risk patient. First, to evaluate the separability of the data, an unsupervised algorithm was implemented. After determining the optimal number of groups (i.e., two classes), supervised machine-learning algorithms (SVM and MLP) were utilized. When we compared the outcomes of these approaches, it was clear that the SVM model delivered the highest accuracy for classifying LBP patients with a maximum accurate level of 75.4% (Table 2). Finally, using the SVM approach, other feature sets such as FT16, Wii, and TSK (see Table 1) were added to the feature vector. Accuracy-related findings for the full factorial combination of the feature sets confirmed that the IMU signal (FS feature set) remains the best feature for identifying LBP patients (Table 3).

As noted earlier, we had originally envisioned that this study would include three distinct categories of LBP sufferers: low, medium, and high risk. However, the unsupervised machine learning approach we employed determined the optimal number of classification groups to be two. Moreover, supervised machine learning for three-class classification resulted in moderate accuracy. Thus, based on our earlier discussion on the importance of discriminating the high-risk LBP patients, it was decided to complete the analysis for the 2-class classification. This decision was bolstered by the fact that, according to clinical treatment protocols associated with the three risk groups, there are no significant differences in the treatment of low and medium risk LBP patients as identified by the STarT questionnaire [19]. Hence, discriminating between high vs. low-medium risk patients would still be valuable and fulfill the objectives of this investigation. Interestingly, outcomes from the SVM model also demonstrated the substantially better discriminative capability of the motion-sensor signal to distinguish patients with high risk vs. low-medium risk for LBP (Table 2).

It is noteworthy to mention that a recent study was successful in discriminating between healthy subjects and lower back pain sufferers based on trunk kinematics [21]. However, differences in trunk kinematic data between a healthy cohort and a patient group are likely to be more significant in comparison to differences within groups of LBP patients at variable risk levels. Moreover, partitioning the subjects into healthy and LBP groups could be performed more accurately in comparison to subgrouping LBP patients into three risk levels. It should also be noted that there is a gray zone for the reference method—for this study the STarT questionnaire—in terms of the appropriate group to which to assign a patient. For example, according to the STarT questionnaire, if a patient’s score was less than 4, s/he would be assigned to the low-risk group. The question, however, is the extent to which questionnaire can successfully assign those on the cusp to the proper group, whether low- or medium-risk. Hence, the uncertainty in labeling the “cusp patients” to one group or another could also introduce some inaccuracy in results obtained from the motion-based classification. Consequently, less accuracy was actually anticipated in the partitioning of the LBP patients compared to discriminating between healthy vs. LBP group.

Since the accuracy of K-means clustering was not sufficiently high, we concluded that there was no clear data boundary between the high and low-medium risk groups. This analysis could signal the necessity to implement supervised algorithms to develop models with higher accuracy. Since there was no similar study designed to classify LBP patients such that there are different treatment approaches for each class, we then we compared our model to other machine-learning-based models that were developed for clinical decision making [45–47]. The obtained accuracy for the best machine learning model (i.e., the SVM algorithms) in this study was in the range of clinical decision-making models described in the literature that were based on signals from IMU or body-worn sensors (i.e., <80%) [45–48]. Compared to the SVM model, the MLP approach achieved a lower accuracy of ~60%, which could be related to the fairly small sample size in this study (94 for 3-class and about 65 for 2-class). In comparison, other analogous studies have included a greater number of observations for each class [49, 50], and in particular, for implementing a neural network approach [51].

The accuracy of the SVM model was investigated in connection with the different feature sets. First we confirmed that COP data for the Wii feature were inadequate for discriminating between high vs. low-medium risk LBP patients (accuracy of ~40%). However, the ADT feature set (which included psychological information such as anxiety and depression) achieved better results compared to the Wii feature set. The FT16 feature set, which had 16 statistical measures of the IMU signal, led to better accuracy (~67%) and specificity (~76%) compared to ADT. However, they demonstrated roughly the same capability in identifying high-risk subjects (sensitivity of ~56% and ~58% for ADT and FT16, respectively). Overall, the kinematic feature set of FS was by far better than the others individually, or in combination with the FS signal itself (see Table 3).

Currently, LBP patients have to go to clinics to complete the STarT questionnaire in order to determine their risk status. Additionally, a qualified care provider must analyze the results of the questionnaire in order to assign the patient to a treatment path. Accordingly, the findings from this investigation could facilitate the development of an objective tool to provide insights on the risk/level of LBP without requiring people to visit their medical provider—at least initially. Another important aspect of this study is that the presented approach only utilized the data from one motion sensor during trunk flexion/extension in the sagittal plane. This setup was designed in order to make the data collection as simple and quick as possible without the help of an expert. Hence, it could be implemented in an application and installed on most of the current smartphones, which as we know are already ubiquitous and are expected to grow in sophistication [52]. Thus, by embedding the developed algorithm of this study on a smartphone application, assessing the risk factors for LBP patients would become easier and more accessible for patients in need.

This study features several limitations that should be addressed in future investigations. The first limitation is that all of the participants were male. Therefore, utilizing the developed models for the female population should be undertaken with a certain degree of caution. A future study should include a gender-balanced population to address this shortcoming. A second limitation pertains to the moderate sample-size. Accordingly, a follow-on investigation should recruit a larger sample size, which could enhance the generalizability of findings and the accuracy of machine learning algorithms. Investigating patient clustering during the treatment process also represents an interesting opportunity for future research. Specifically, a study could be designed and conducted that would compare the impact of treatment processes to patient kinematics and/or self-reported questionnaire data.

## Conclusion

In our study, we assessed the capability of motion-capture sensors as an objective measurement tool to be implemented as a clinical decision-making scale instead of STarT questionnaire, which is a subjective scale for classifying LBP patients. Our results showed that machine learning methods, especially SVM, can distinguish high vs. low-medium risk LBP patients with adequate accuracy (>75%). The outcome of this exploratory study could be used to develop an objective tool capable of classifying the patients to better assign them to the proper treatment path. Finally, the findings from this investigation could facilitate the development of healthcare prognosis tools that would help to assess low back pain sufferers with greater ease, more objectively, and at lower cost in comparison to current clinic-based methods.

